# Genetic Information Relationship Network (GIRN): A Force-directed graphing tool for gene expression analysis

**DOI:** 10.1101/079590

**Authors:** Gaelle Rondeau, John Welsh

## Abstract

We present a web-based tool, Genetic Information Relationship Network (GIRN), for mapping genes to a global protein interaction network for humans, and for further annotating this map with gene ontology, pathway, disease, and user-generated terms. These annotations can be adjusted according to enrichment within a subset of genes. Additionally, drugs that interact with genes in the subset can be added to the graph. The maps are force-directed graphs in which highly connected nodes tend toward the center, and highly interconnected nodes tend toward each other. Icons of different shapes and colors are employed to indicate whether the node represents a protein, a gene, or one of the various types of annotation. Coordinated interaction of genes and functional inference can be identified by visual inspection. Each node in the graph is associated with a menu of links to external data sources. Collectively, these tools provide an efficient portal to gene-associated public data for any group of genes specified by the user. The site can be found at www.voxvill.org/relnet.

## Introduction

Microarrays and RNA-seq are straightforward technologies to identify genes having expression behavior that correlates with other variables. However, it remains challenging to create meaningful narratives describing the roles played by different genes in complex biological systems. Such meaningful narratives might assume the form of review-like descriptions of the behavior of a gene, and in computer science might be formulated as object models. An important objective of constructing accurate narratives describing gene function from large data collections is to lead us to a deeper understanding of the basis of biological phenomena, and in practical terms, to derive actionable knowledge, such as the identities of genes that might be targeted to treat diseases. The challenge stems largely from the combinatorial complexity of biology, wherein most genes are involved in more than one process, most processes involve multiple genes, and ultimately no process is fully isolated from other processes. The vast combinatorial complexity of biology suggests that computational methods will be helpful. Substantial effort has been directed toward building structured, machine-readable biological information and algorithms to manipulate that information to automatically or semi-automatically derive useful inferences. These databases often take different kinds of information into account. For example, The Gene Ontology (GO) Consortium (1, 2) have grouped genes according to biological process, cellular location, and molecular function, whereas Reactome (3) provides a powerful interface to explore gene expression data in the larger context of GO, but with an emphasis on pathways. STRING uses detailed information about protein interactions over many species (4), whereas HTRIdb contains experimentally verified human transcriptional regulation interactions (5), DGIdb is a database that keeps track of drug-gene interactions (6), and DisGeNET(7–10) relates genes to diseases. Consensus Path DataBase (11) leverages the composite data of >30 databases concerned with various aspects of gene function. The NDEx (Network Data Exchange) Project (12) provides an open-source framework for community-wide involvement in the advancement of biological network information.

Most strategies for analysis of differential gene expression rank genes according to one or more criteria and place them in tabular form, which usually serves as starting point for both manual and computer-assisted functional analysis. For example, the top 100 genes that respond to an experimental drug can be examined for enrichment of GO, pathway, disease, or other functionally relevant annotations using a variety of web-based tools, presented in tabular form. Several of these tools also employ graphical network-based concepts, e.g. BioGRID (13), Consensus Path DataBase (11), Pathway Commons (14), Reactome (3, 15), STRING (4), PathwayNet (16), and others, or enable network display on third party software (e.g. Cytoscape, VisANT), such as NetPath (17). Pathway enrichment is typically achieved using a binomial proportions test (18). Another approach, weighted gene co-expression network analysis (WGCNA), organizes genes in a network according to adjacency functions based on co-expression (19).

Network graphs are generally useful as data summaries when they allow one to quickly identify interacting entities, clusters of interacting entities, intermediates between clusters, and hubs. Here, we present a website that takes a list of genes as input, identifies and displays the local interaction sub-network as a force-directed graph, with genes, interaction type, and annotation terms as nodes. Physical interactions between proteins, between transcription factors and the genes they regulate, and between proteins and small molecules, such as drugs, are all displayed in a single composite network graph. In addition, diverse kinds of annotation information can be displayed using the same format without modification.

## Results

### Graph conventions

In this website, genes, gene ontology terms, pathway descriptors, drugs and diseases are all represented as nodes in a force-directed graph, and their interactions are represented as edges. Analysis is initiated by entering a set of genes or choosing from a variety of pre-loaded gene lists, and several analysis options are discussed in the sections that follow. In these graphs, different icons are used to signify the database to which each node belongs. Each edge is interrupted with a centrally located icon signifying type of interaction between nodes. All nodes are clickable links: left double-clicking opens a link to one or more relevant references in a new tab, and right clicking opens a drop-down box to other links to relevant information, or to websites likely to contain relevant information, depending on the type of node. The network image can be zoomed and repositioned via simple key board and/or mouse actions. Combined, these features allow for convenient and rapid review of structured knowledge surrounding the sub-network of interacting genes subtended by a gene list.

### Access to lists of genes associated with gene ontology terms, pathways, diseases, and drugs

The site is primarily designed for the purpose of inferring the relationships of a group of genes supplied by the user to known biology. However, one is often interested in exploring the genes associated with established gene ontology, pathways, diseases, and drugs that comprise the knowledge base upon which such analysis depends. As a convenience, the site provides a convenient text-based boolean search tool to identify genes associated with GO terms, pathways, diseases, and drugs, although sites dedicated to these individual areas, such as those maintained by The Gene Ontology Consortium (1, 2) (www.geneontology.org), Reactome (3, 15) (http://www.reactome.org) , DGIdb (6) (http://dgidb.genome.wustl.edu), DisGeNET ((http://www.disgenet.org) (7, 9), and others, often provide more sophisticated search capabilities. Those sites may be useful to identify terms of interest or gene lists. Either route allows one to choose genes associated with various classifications, and these can be studied in isolation or in combination with other sets of genes.

### Identification of genes that participate in protein-protein and transcription factor-gene interactions with genes in a list

Methods such as microarrays or RNA-seq are used to identify lists of genes associated in some way with a phenotype or other observable, and there is interest in drawing inferences from such a list of genes and other sources of structured information. An assumption is that genes in the list are unified by some underlying rule, and a common goal is to identify the rule. One generic approach is to display the list of genes together with genes that can be inferred as “neighbors” in the global network of protein-protein and protein-gene interactions. Analysis is initiated by pasting a list of gene names, specifically official gene symbols (HUGO Gene Nomenclature Committee, HGNC; http://www.genenames.org (20)), into a text box and selecting a z-score. At this level of analysis, low z-scores display most or all genes that are connected to the genes in the list, and this is often of interest because the connection of even a single gene in the list to some gene that is important in the global network may be of interest. It is possible to exceed available server resources when large lists of genes are returned, in which case output is limited to genes with the highest z-scores. Higher z-scores restrict the network output to genes that are more highly interconnected than one would expect based on random chance. Inclusion of such genes occurs when two or more of the genes in the input list are connected, and this behavior is expected for genes that exist as part of a complex, or that share a pathway. In these instances, the z-score may be used as a means of ranking competing hypotheses regarding which pathways or functions are implicated by a list of genes. It will be generally useful to examine the network at various z-score levels. Figures 1a-c illustrate the effects of using different z-scores to define the network.

**Figure 1a-c.**
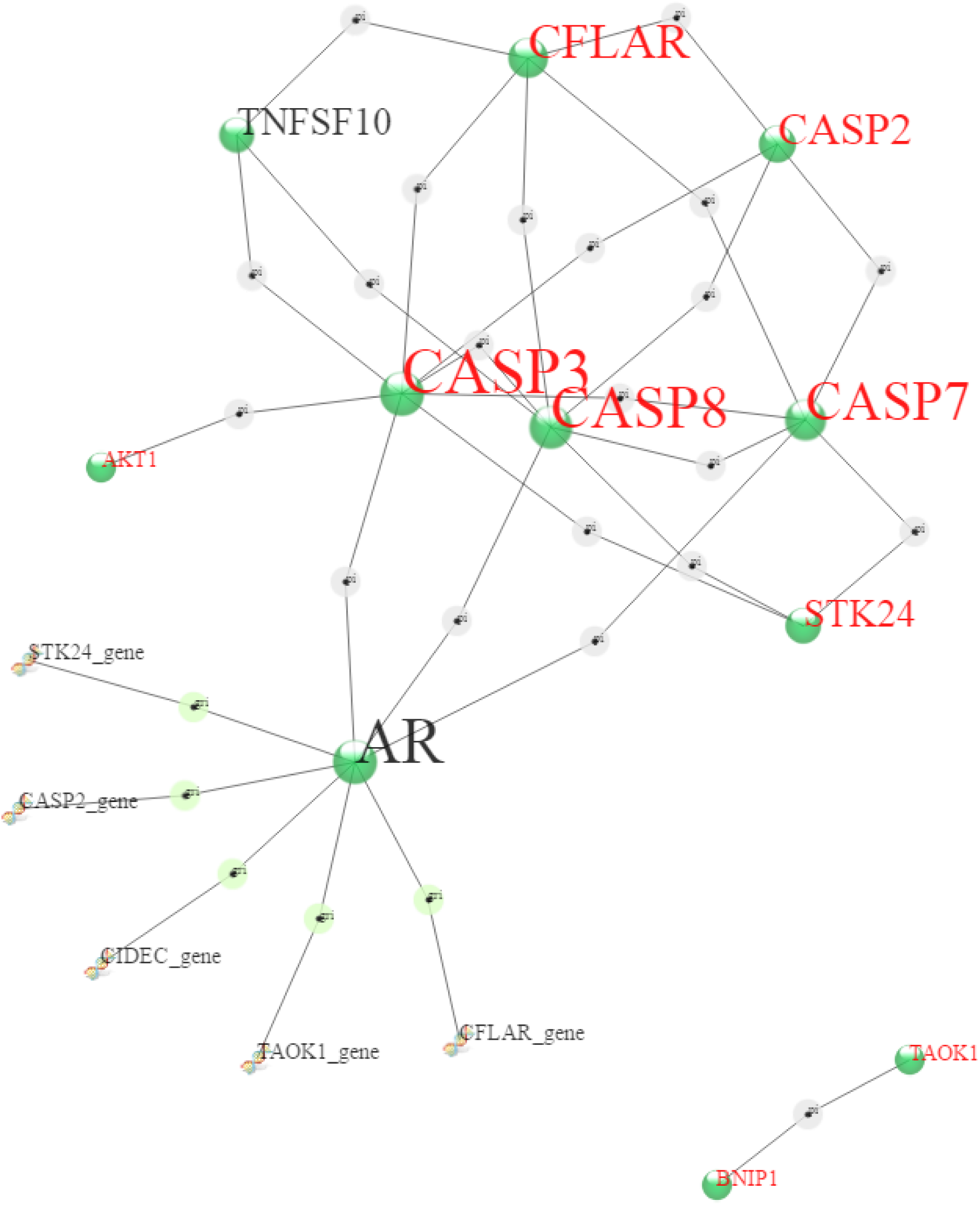
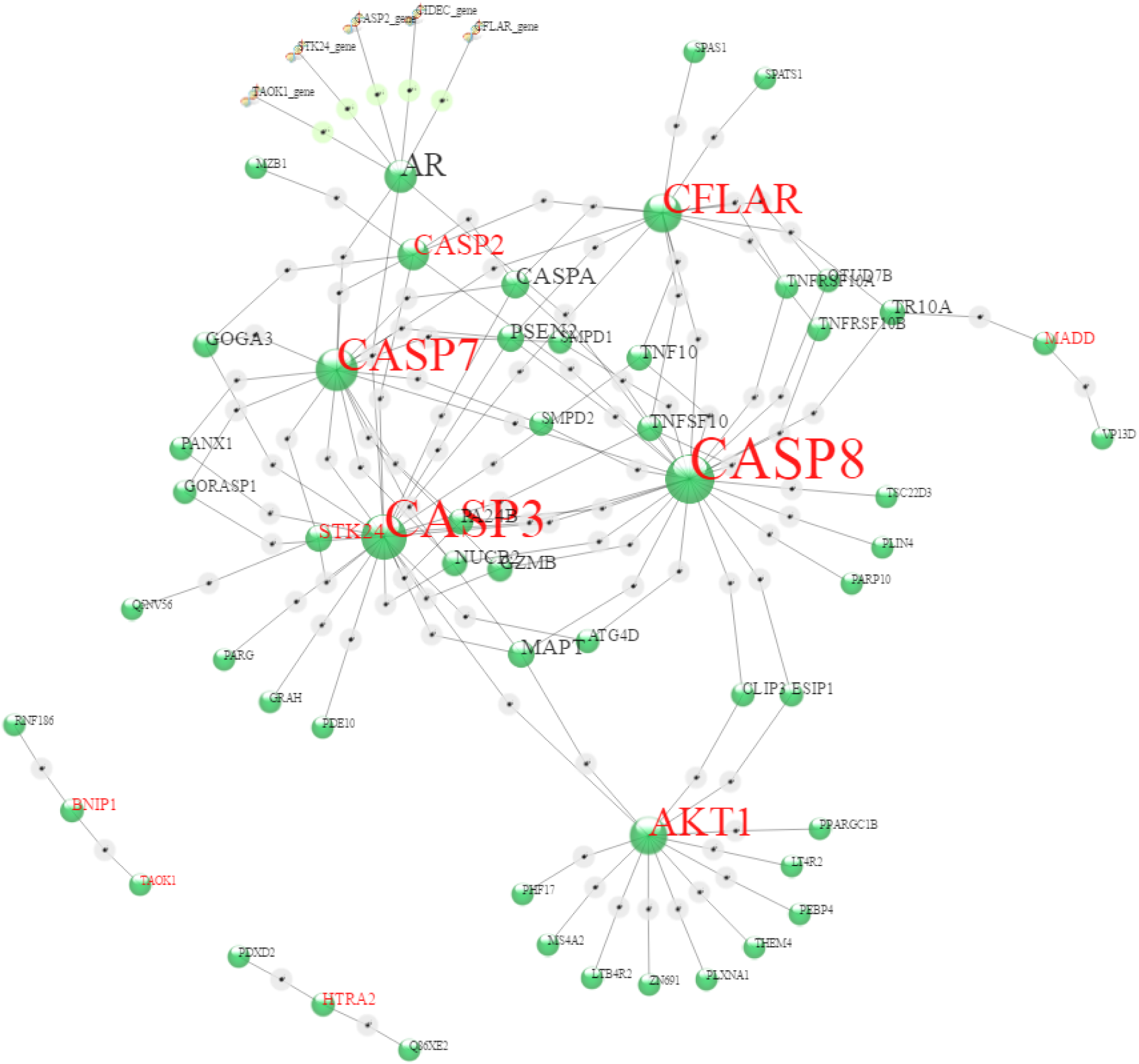
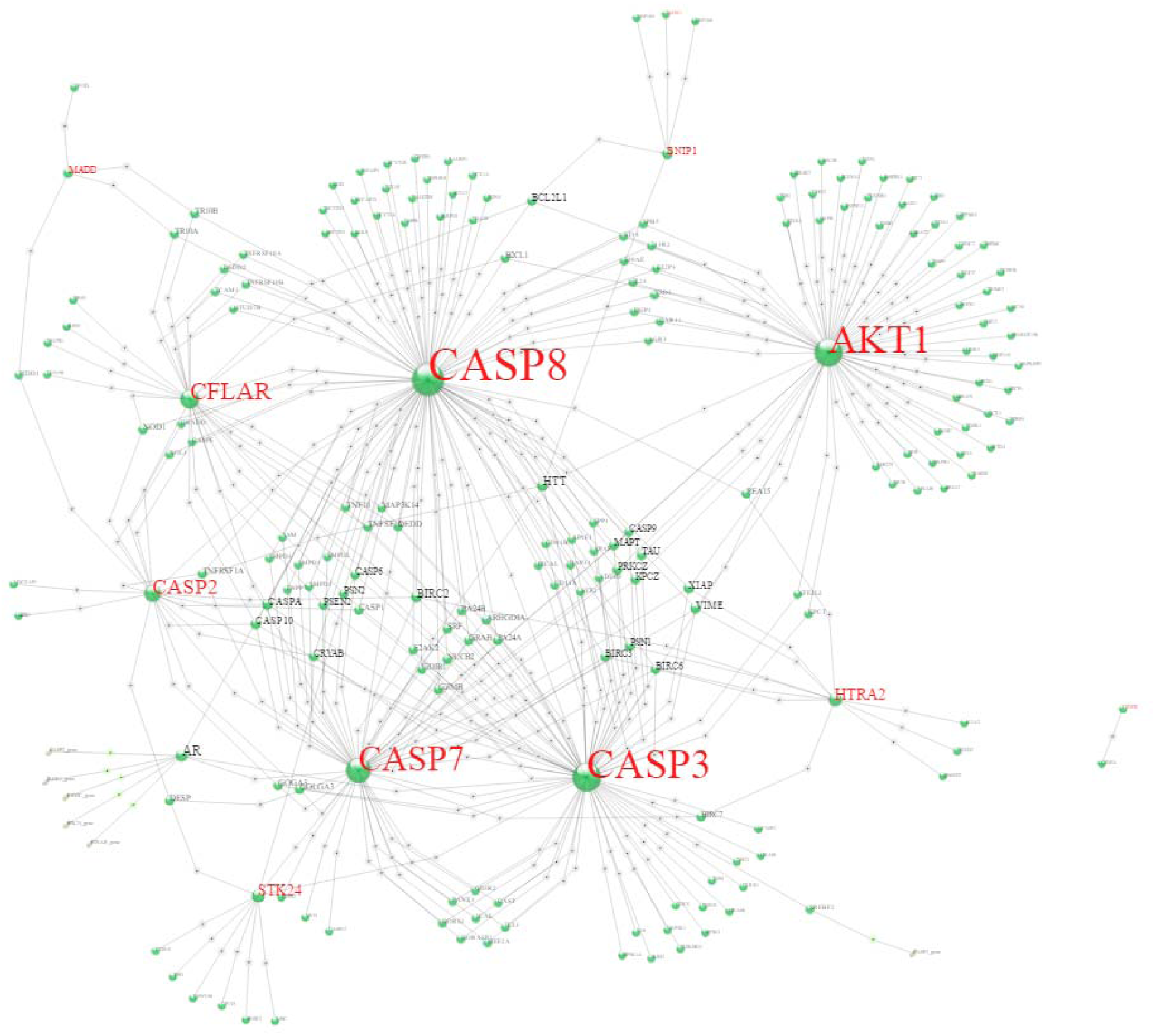
The size of the network can be adjusted by changing the z-score. Gene ontology terms for “GO:0097194 execution phase of apoptosis” are used in this example. (a) z-score = 400, (b) z-score = 200, (c) z-score = 100.

Force directed graphs are appealing because they self-organize. These graphs tend to self-organize in such a manner that the more highly connected nodes are located toward the middle. As an added visual enhancement, the font size of the text label scales according to the number of connections, which has the effect of drawing the eye to the more highly interconnected nodes. Figure 2 shows a self-organized force directed graph as it self-organizes. All node positions can be fixed or released, by clicking or shift-clicking, respectively. A node can be selected as a focus (Figure 3a-b). All nodes are clickable links, with right clicking opening a drop-down menu to other sites containing gene-centric, ontology, pathway, gene set, drug, or disease information, depending on the class of the node (Figure 4). Left double-clicking opens a new tab with type-dependant additional information.

**Figure 2.**
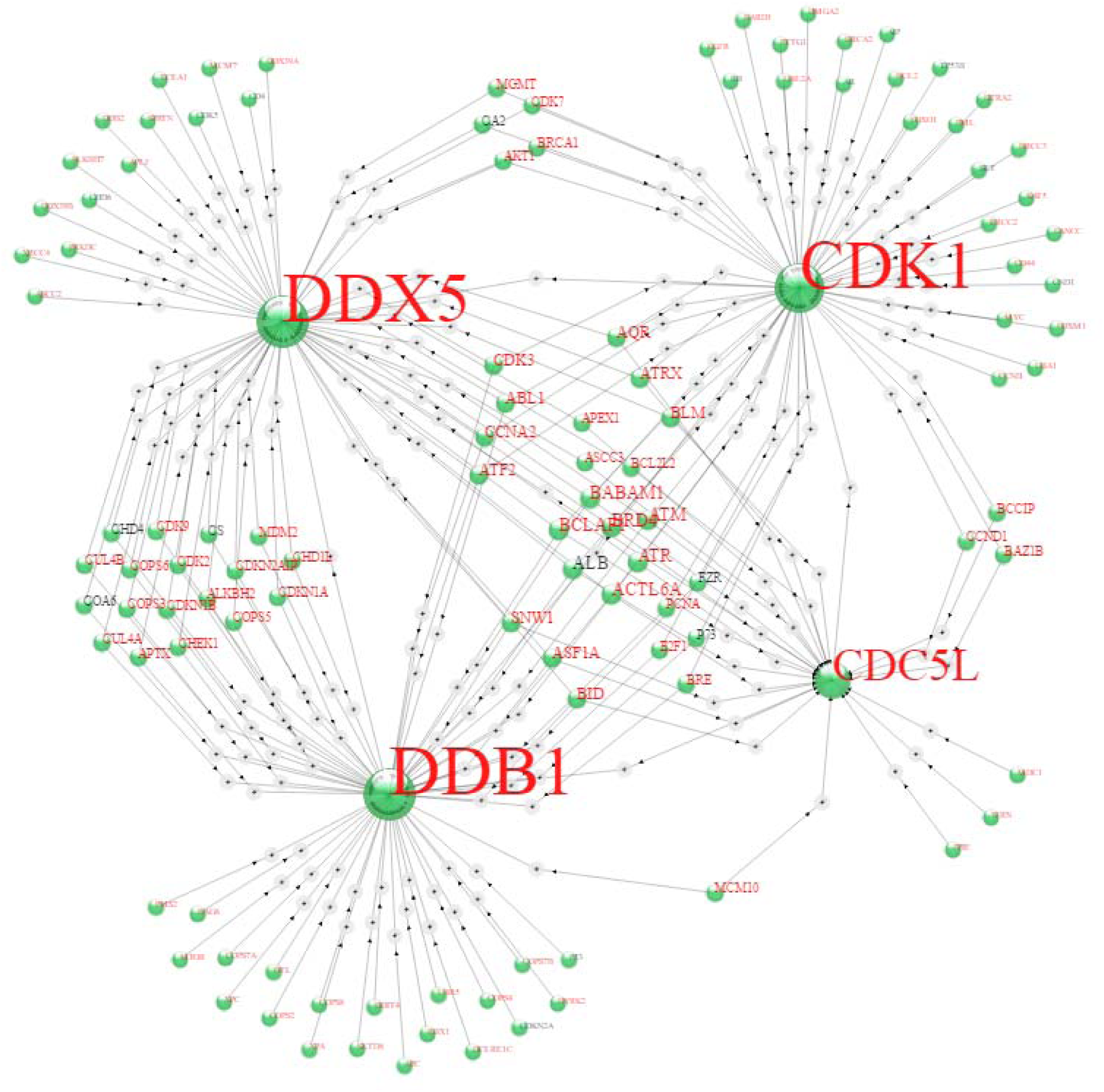
Networks self-organize. Genes associated with all GO terms matching “DNA damage || DNA repair” (the double pipe indicates “OR”). Max genes set to 800 and z-score to 300. UBC was omitted for clarity (by placing “UBC” in the “Omit” box).

**Figure 3.**
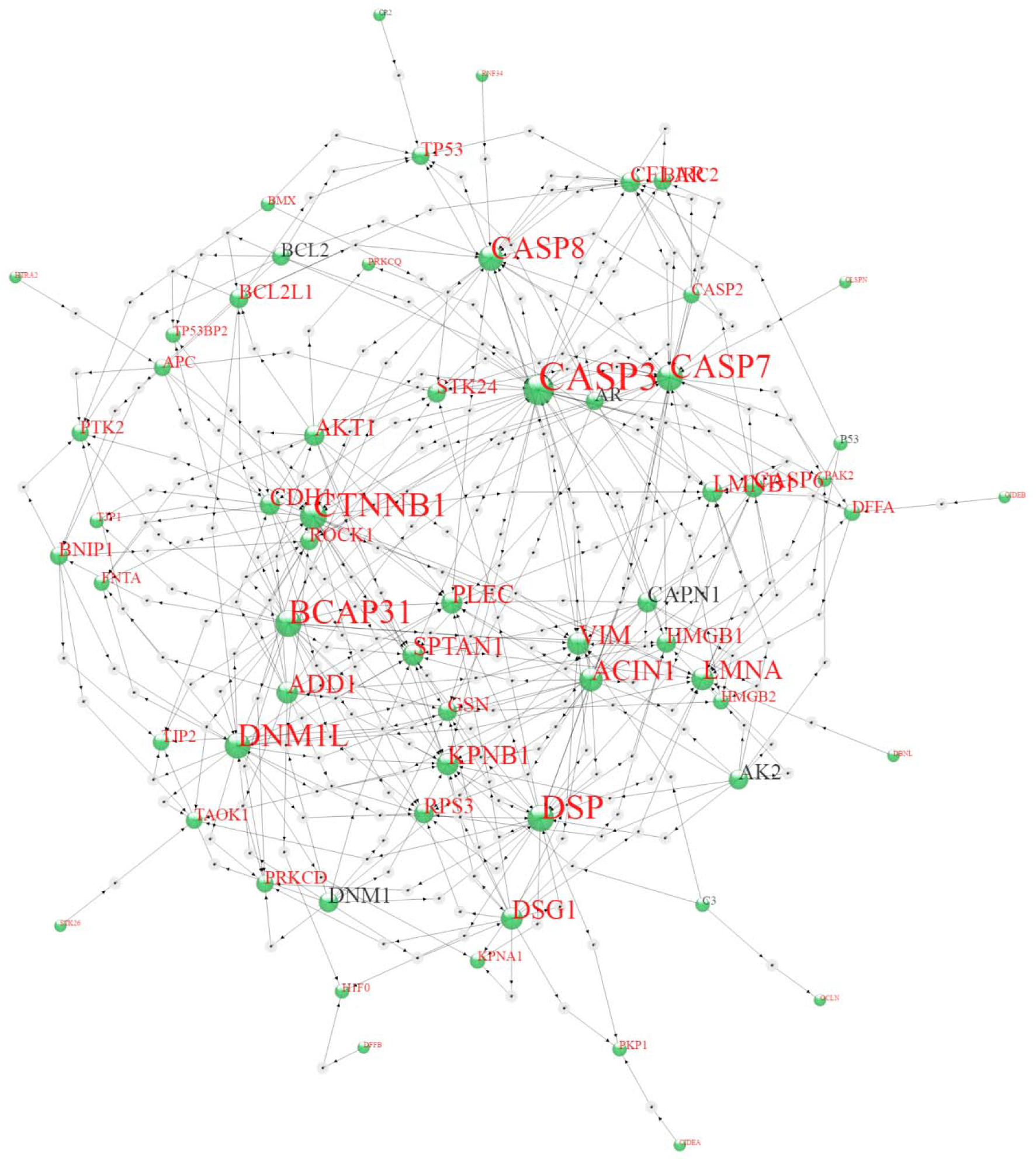
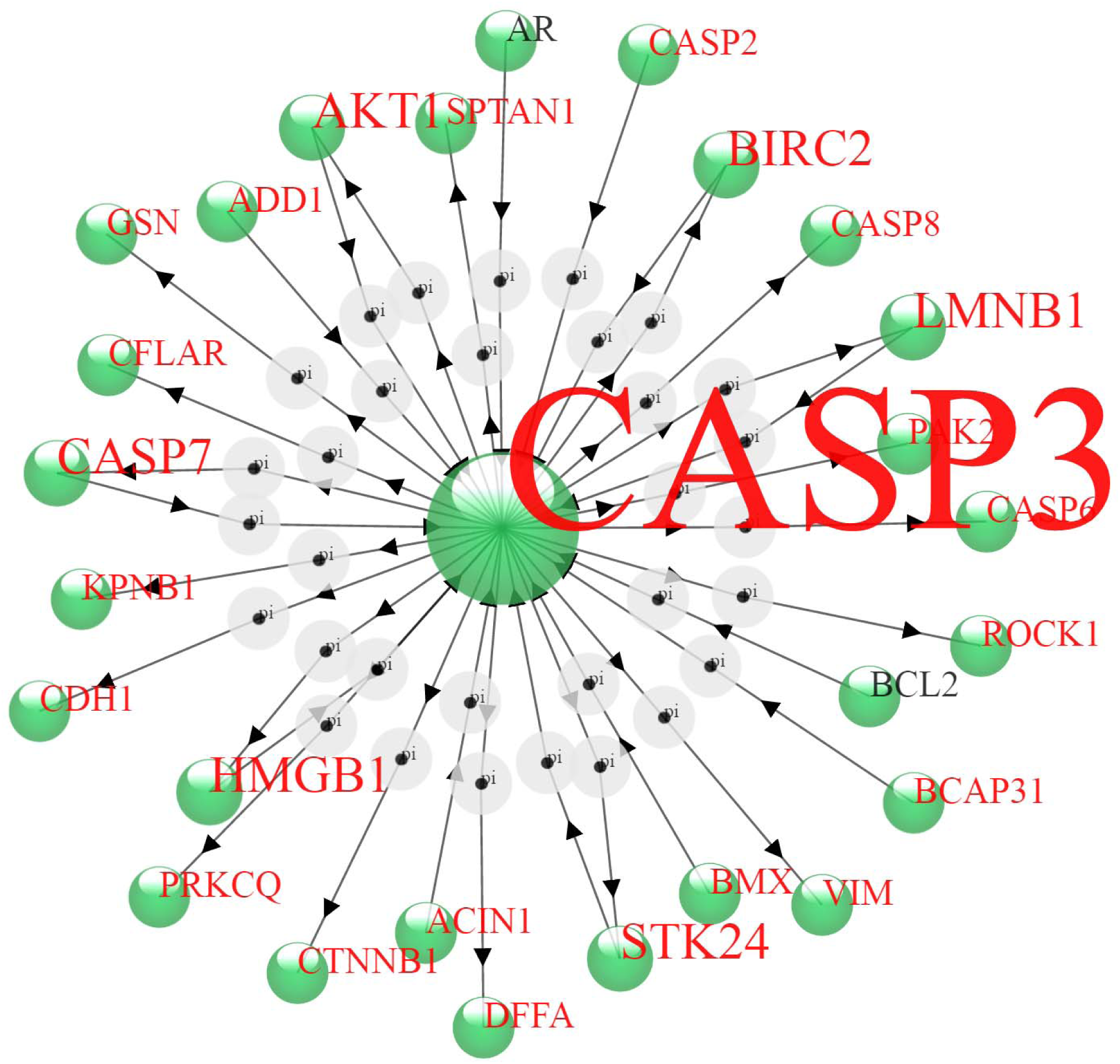
Gene ontology terms containing the word “apoptosis”. (a) includes the subnet at z-score = 400, (b) same as (a) except focus has been placed on genes within one step of CASP3.

**Figure 4.**
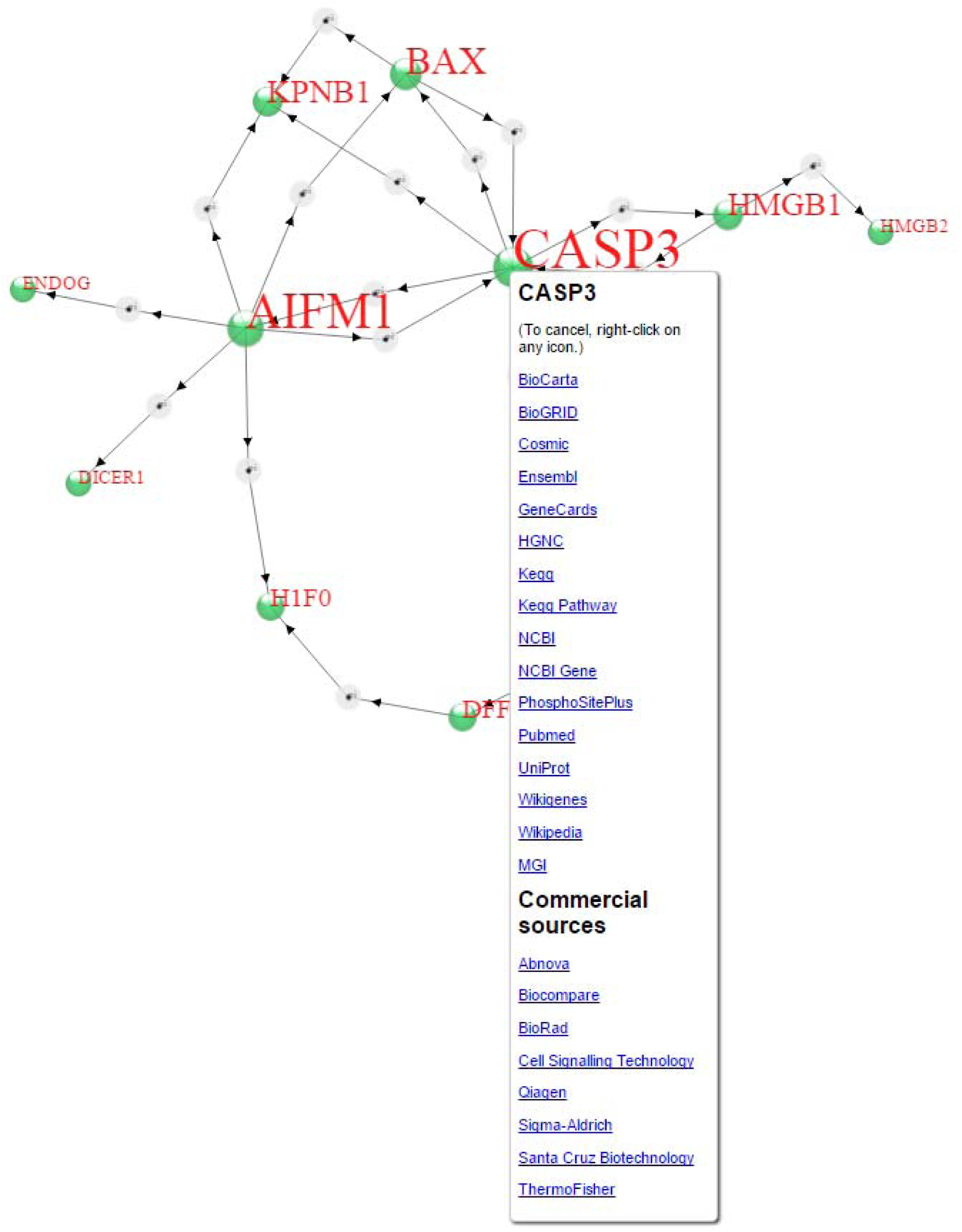
Right-click menu, which displays links to additional information.

The protein-protein interaction network used by this website is a composite from Kamburov et al. (11, 21–23) in Consensus Path Database (http://consensuspathdb.org), from Cerami et al.(14) in Pathway Commons (http://www.pathwaycommons.org) and the transcription factor-gene interaction network is from experimentally verified human transcriptional regulation interactions compiled by Bovolenta et al. (5) in HTRIdb (http://www.lbbc.ibb.unesp.br/htri). These, in turn, comprise composite databases from many other sources, which are referenced accordingly in the above literature citations. Some of these databases are subject to ongoing curation, whereas others are static and some no longer available at their original web addresses. We provide a link to “Biomedical Interaction Databases” on the main analysis page. The z-scores and the p-values discussed herein are based on data that may not satisfy distribution assumptions, and therefore may not accurately reflect statistical confidence. In addition, the completeness of the global protein-protein interaction network is difficult to assess, a priori. As these z-scores and p-values are intended to rank genes or attributes according to the chances that they are part of an inferred sub-network, there are bound to be errors in ranking. Overall, however, we expect rankings to be approximately correct. Also, it should be kept in mind that a change in the expression of a single gene can radically alter cellular behavior; therefore, the rationale justifying the use of “enrichment” as a strategy to rank genes or attributes according to importance should be applied with care.

In these networks, proteins are indicated by green marble icons, whereas genes, where explicitly invoked, are indicated by alpha-helix icons. An objective is to develop a set of icons for various categories of annotations, such that association between icon and annotation category becomes second nature, perhaps with a little practice.

A protein product does not generally interact with the gene that encodes it. Nevertheless, such interactions are scored as interactions to allow the network to display the role played by a transcription factor in the ultimate translation of a protein product (Figures 5a,b).

**Figure 5.**
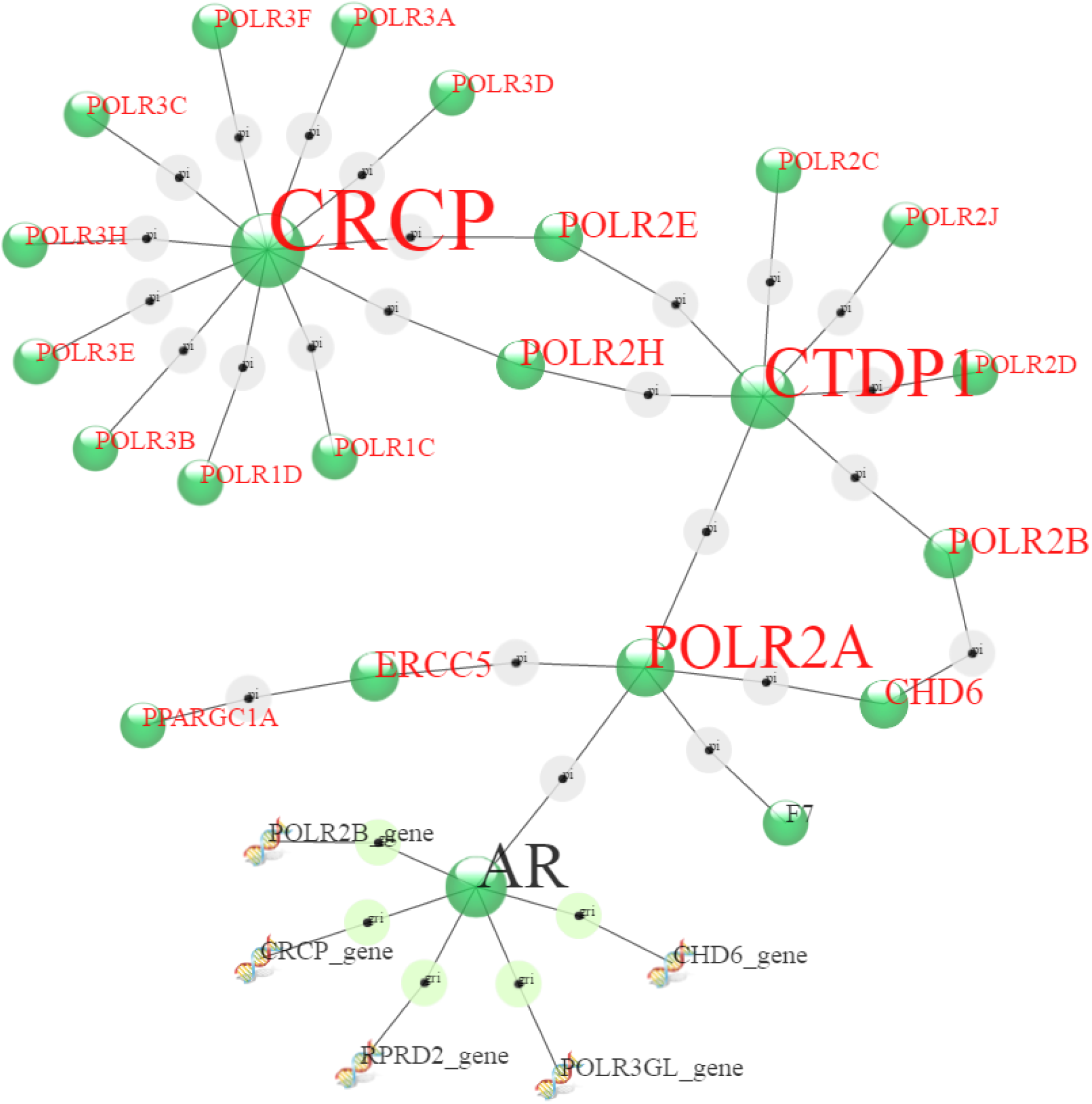
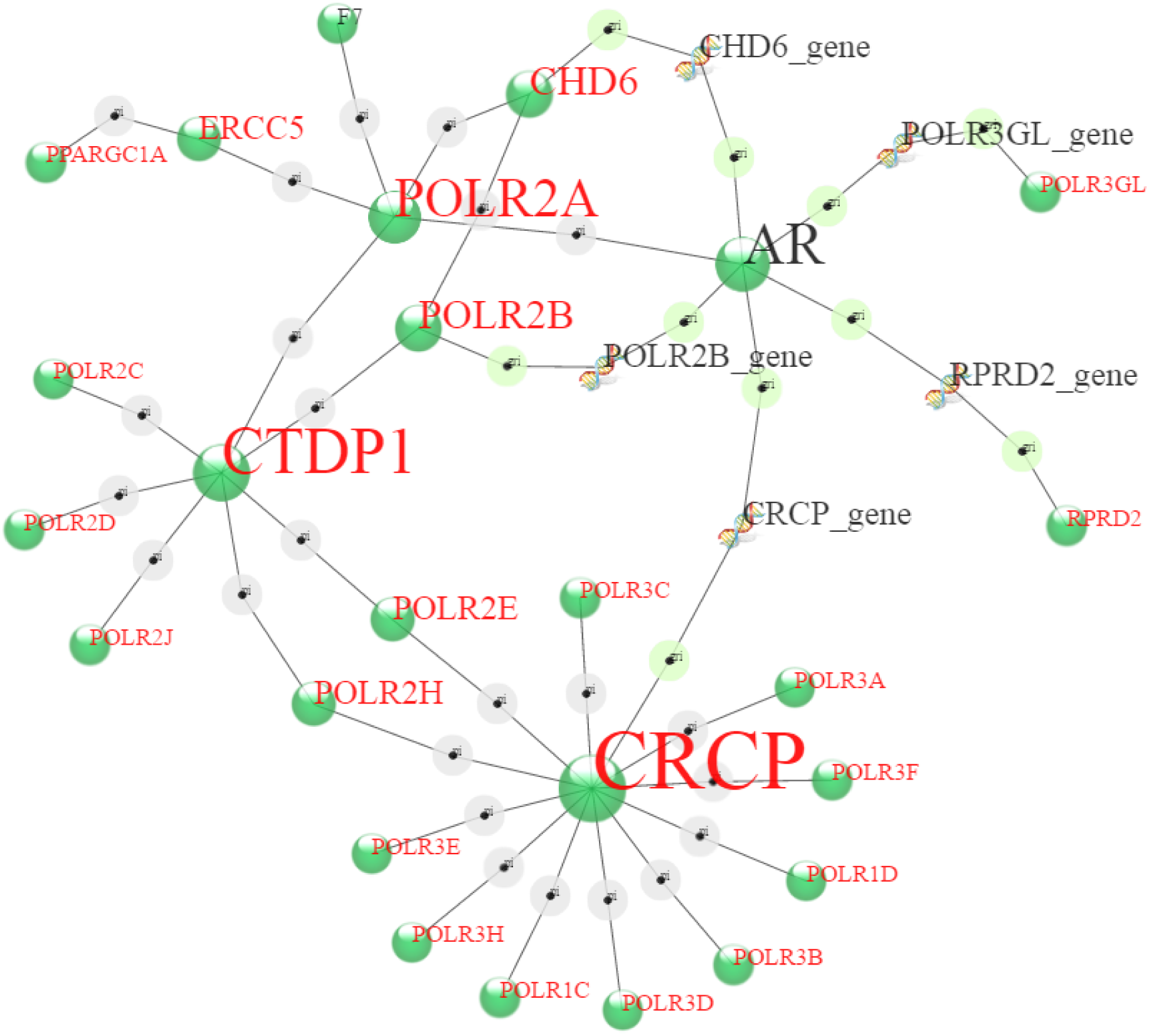
Gene ontology terms containing “DNA-directed RNA polymerase II” (specifically GO:0005665 DNA-directed RNA polymerase II, core complex, GO:0016591 DNA-directed RNA polymerase II, holoenzyme, and GO:000566, DNA-directed RNA polymerase III complex), z-score = 400 (a) without and (b) with edges connecting protein products with their gene names.

### Identification of gene ontology terms associated with genes in a list

There are several predefined lists of genes corresponding to genes associated with gene ontology terms, pathway descriptors, diseases, and genes known to interact with drugs and other small molecules. Two such lists can be selected sequentially and compared. Gene ontology pertains to functional categorization of genes. A common practice is to associate genes in a list with terms in a database in which each gene is mapped to one or more ontology terms. Analysis of enrichment of gene ontology terms informs as to possible changes in phenotype associated with a list of differentially regulated genes. In Figure 6, Gene Ontology Terms are placed in the network as blue, red, or yellow marbles, depending on their class, biological process, molecular function, or cellular component, respectively. Enrichment analysis is performed using Fisher's exact test, which determines the probability of having drawn a sample distribution with an equal or more extreme statistic. In this case, it determines the probability that the several genes presented to the software would contain some number of genes from one or more GO categories. This is a multiple testing environment and the p-values are adjusted using the Bonferoni correction method. A low p-value cutoff allows one to identify ontology terms that are associated with a greater fraction of the genes in the list than would be expected by chance. Ontology terms are represented in the network as nodes, similarly to the node representation of genes and, indeed, all other annotations. An objective of this sort of analysis is to allow the self-organizing properties of the network to organize functional terms close in space to clusters of functionally cooperating genes. Information that cannot be obtained by mere association is the importance of a gene to implementation of the phenotype pointed to by the ontology term; consequently, that a single connection might be important cannot be excluded.

**Figure 6.**
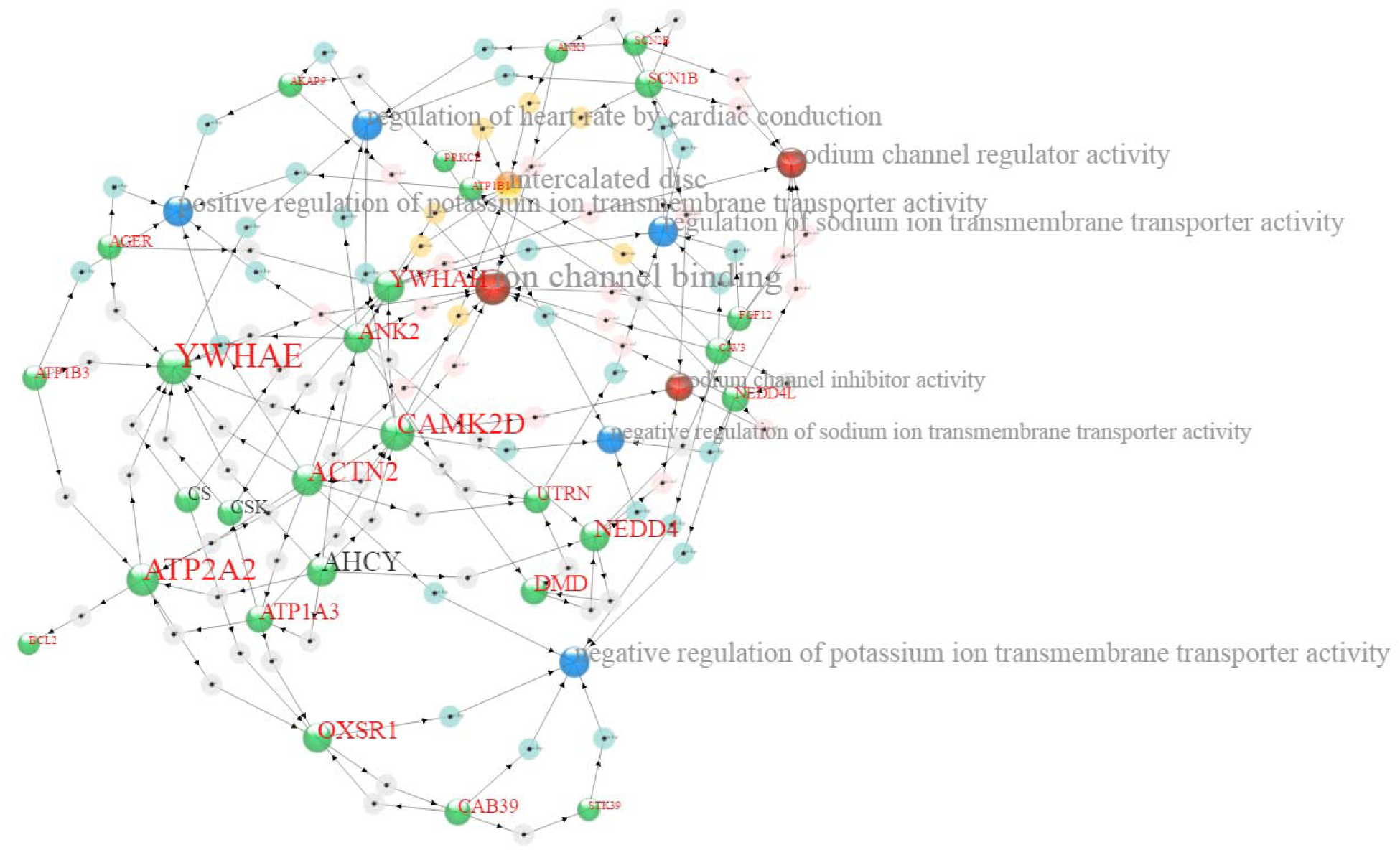
Gene Ontology terms, containing all of the terms “membrane && activity && regulation” (the double ampersand indicates “AND”), z-score = 400, p-value for GO annotations: 1E-6. This shows several cellular component (yellow), molecular function (red), and biological process (blue) terms.

### Identification of pathways associated with a gene list

Pathway data were compiled by Kamburov et al. (11, 21–23) and include data from multiple sources. Genetic Information Relationship Network (GIRN) was used to generate Figure 7, which shows several annotation networks associated with genes from GO terms that contain the string “DNA fragmentation”. A network connecting these genes is presented in unannotated form (Figure 7a), annotated with GO terms (Figure 7b), and annotated with pathway terms (Figure 7c).

**Figure 7a.**
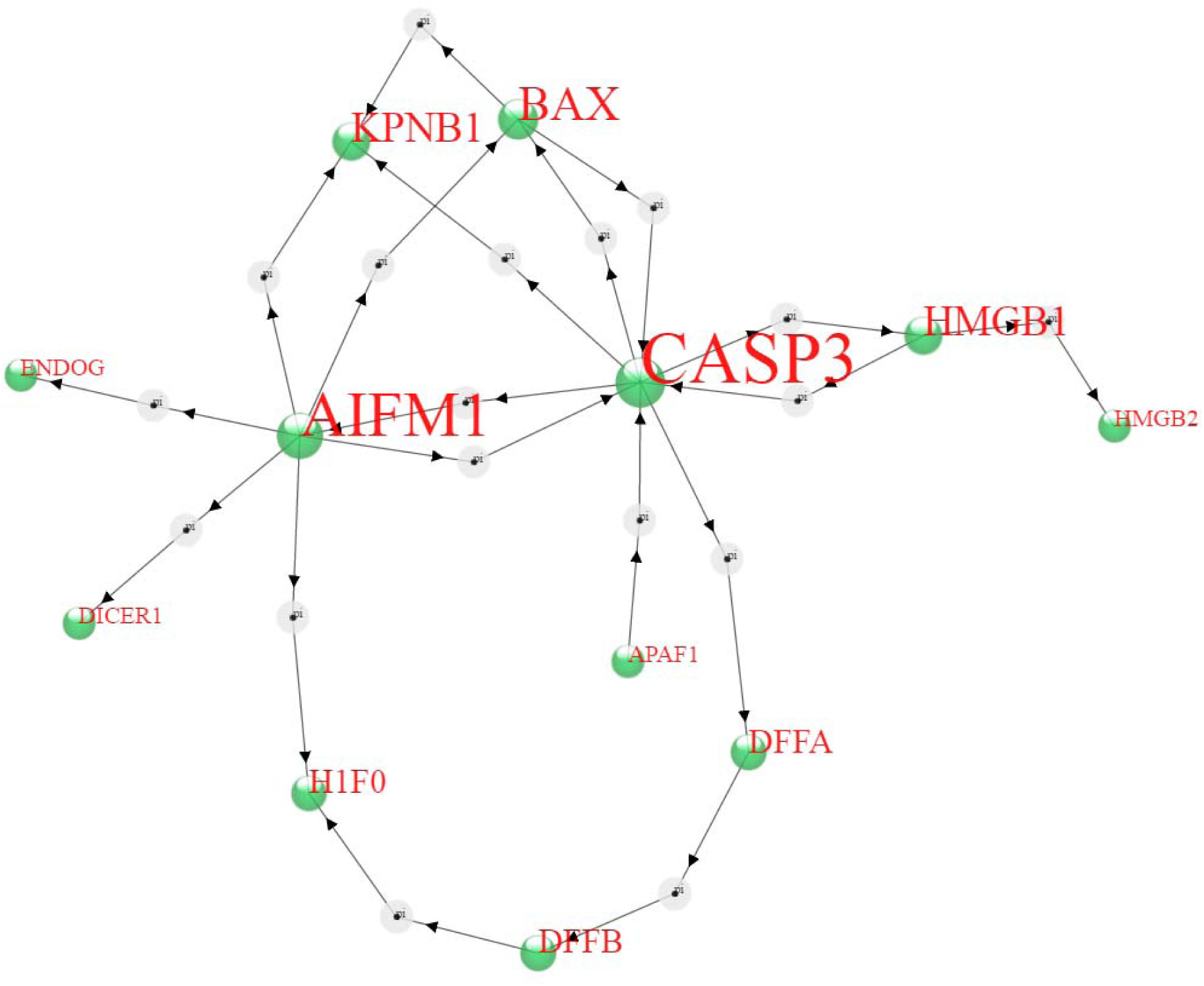
Gene Ontology terms containing “DNA fragmentation”, specifically, GO:0006309 apoptotic DNA fragmentation, GO:1902511 negative regulation of apoptotic DNA fragmentation, GO:1902512 positive regulation of apoptotic DNA fragmentation, and GO:1902510 regulation of apoptotic DNA fragmentation mapped to network using a z-score of 400.

**Figure 7b.**
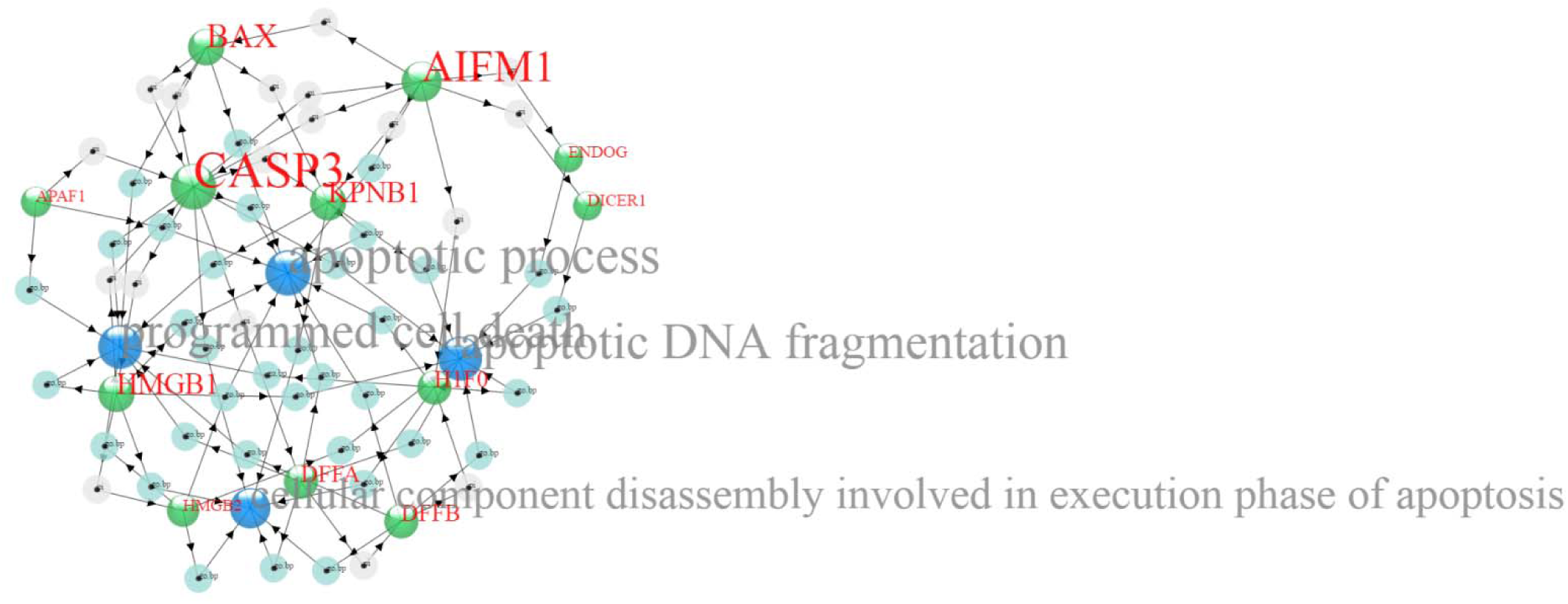
Gene Ontology Same as (a) but annotated with Gene Ontology terms at a p-value = 1E-4.

**Figure 7c.**
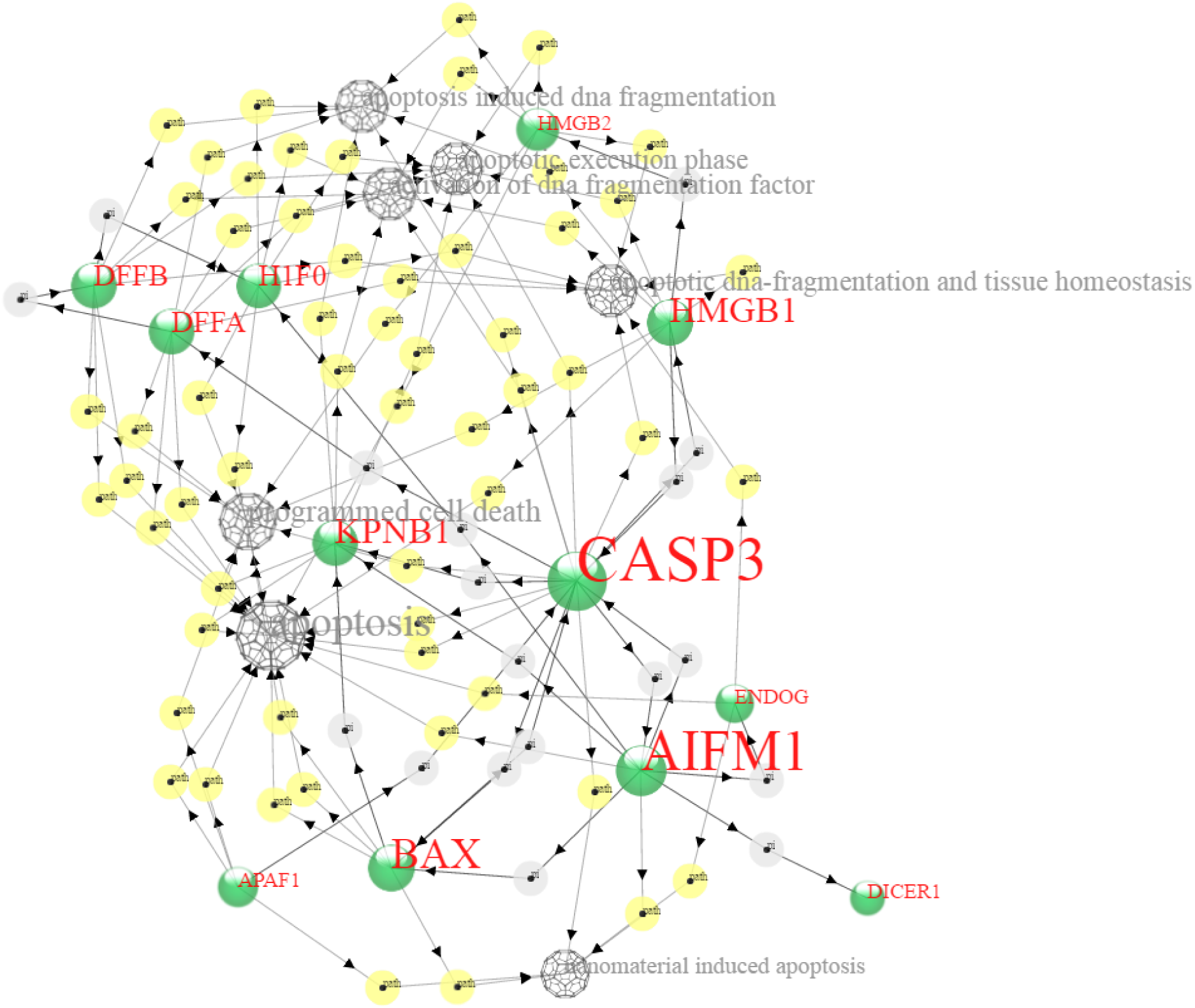
Pathways Same as (a) but annotated with Pathway terms at a p-value = 1E-8.

### Identification of diseases associated with genes in a list

One is often interested in identifying enrichment of genes that are known to be associated with a disease. For example, treatment of cells with a new drug may unexpectedly alter the expression of genes that are associated with one or more diseases, in which case the drug may be a candidate to treat or possibly exacerbate the disease. Disease-gene associations are from DisGeNET (7, 9). Implicated diseases are identified in the same manner as Gene Ontology terms, using Fisher's exact test. Figure 7d shows a network of genes associated with GO terms related to “DNA fragmentation”, and annotated with diseases enriched for genes in this group.

**Figure 7d.**
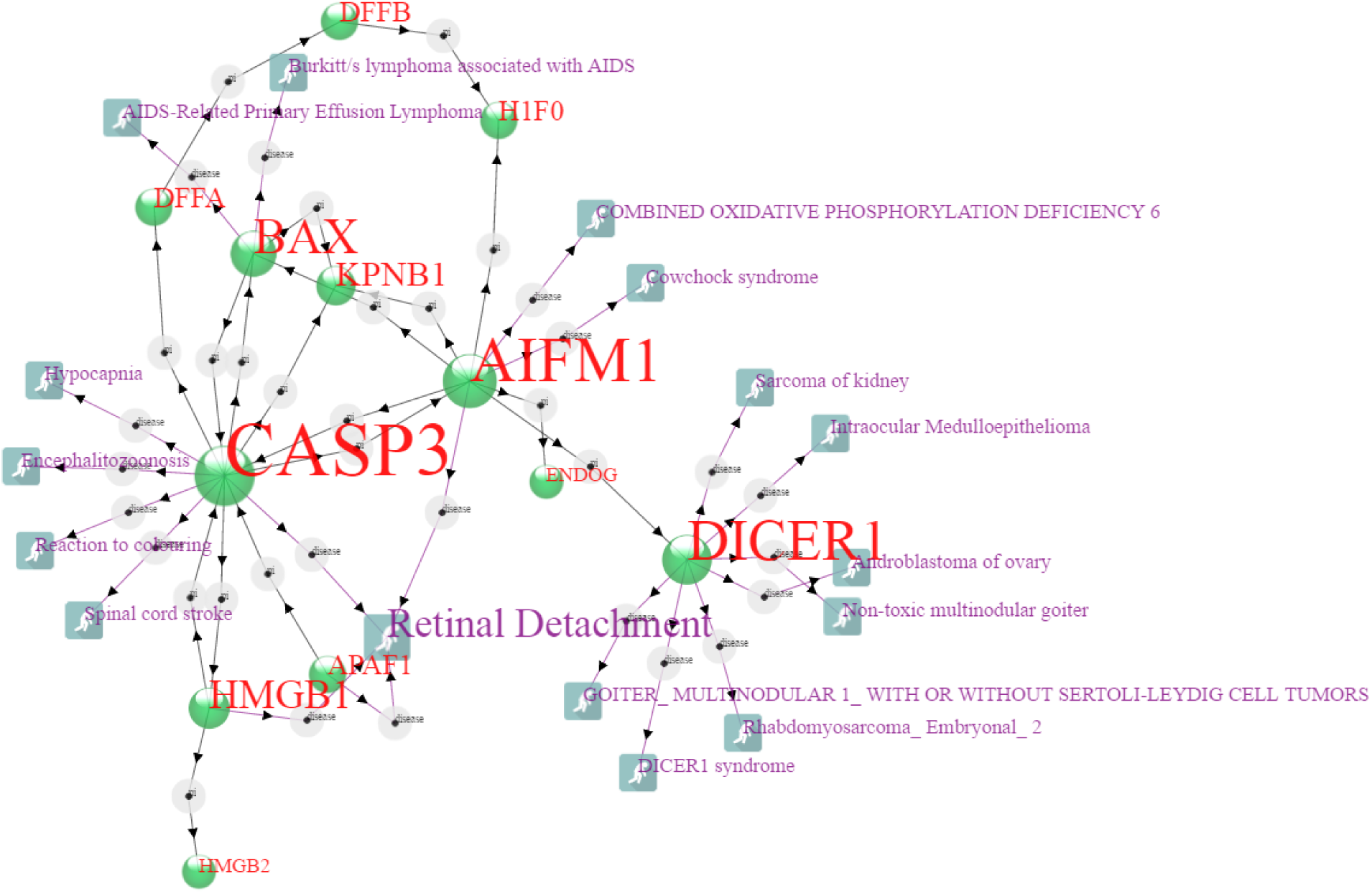
Same as (a) but annotated with Diseases terms at a p-value = 1E-2.

### Identification of drugs that interact with genes in a list

Drug targets. Replace the icon with Rx icon. The drug database is from DGIdb (http://dgidb.genome.wustl.edu). Figure 7e shows the network of genes associated with GO terms related to “DNA fragmentation”, and annotated with drugs that target genes in this group.

**Figure 7e.**
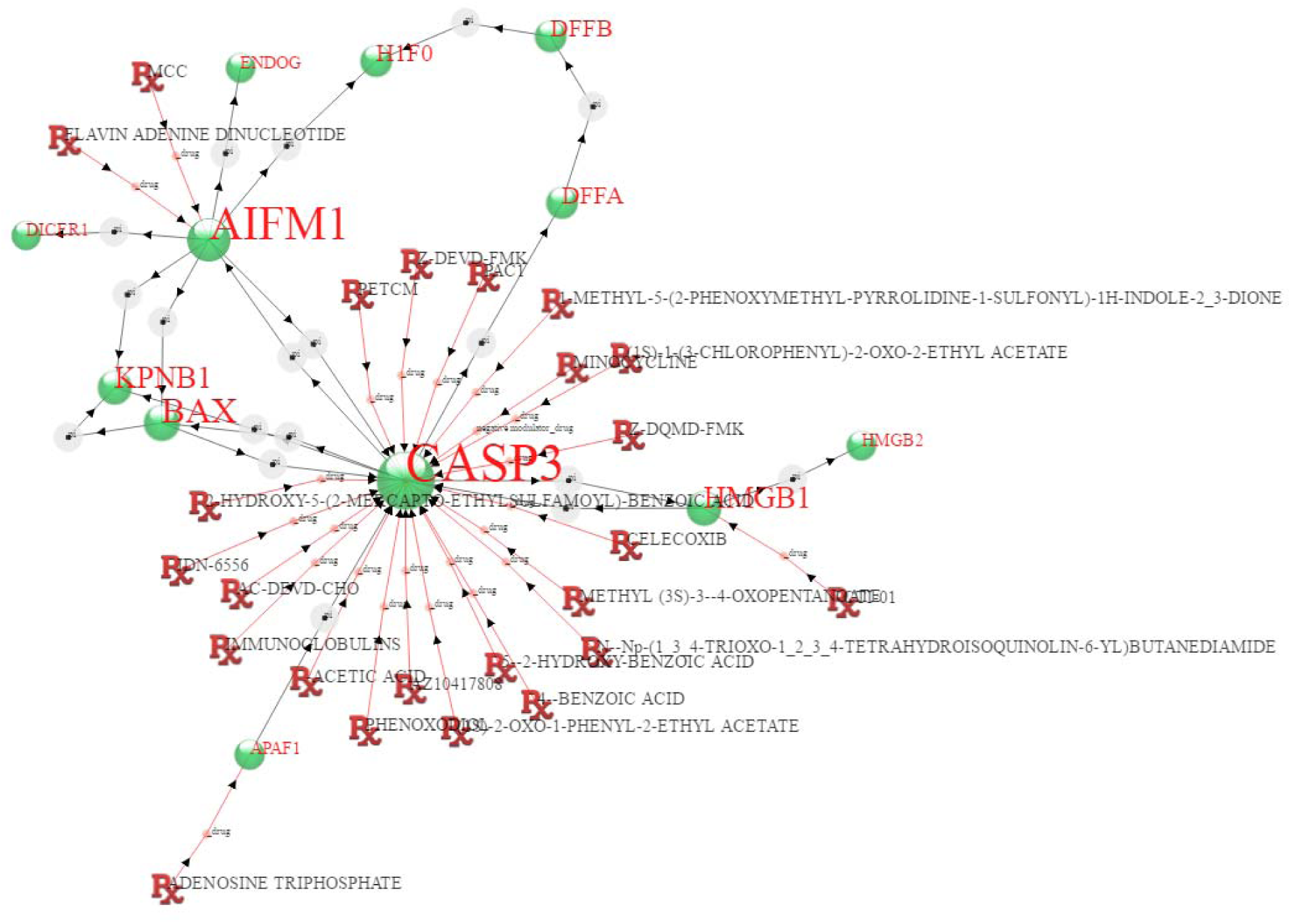
Same as (a) but annotated with drugs.

### Formats allowed

Files using the format from ConsensusPathDB and the Sif format from Pathway Commons, as well as files saved in the comma-delimited format used by GIRN, can be opened using “Choose Files” and are automatically formatted to work in GIRN.

## Discussion

For the most part, biological systems, and malfunctions thereof, are too complex to derive meaningful hypotheses or potential solutions without the aid of substantial, robust, and human-accessible models. However the underlying data are growing very rapidly and methods to extract probable explanations despite humanly intractable complexity will be needed. Machine learning is already used to analyze data in biology, but in these early days of truly vast data, graphical methods may be useful in designing methods to capture information of practical value.

Graphic methods are important in discovery-oriented data analysis and in the communication of results. This is especially true for large data collections, for which graphic representation of summary statistics is standard practice. More elaborate representations of data can be useful when trends can be spotted by eye. For example, heat map representations of gene expression data across dozens or even hundreds of samples are standard practice, and allow scientists to intuitively grasp and convey aspects of similarity and differences between, for example, tumor samples. For a graphic representation to be useful, the data must map to the representation in a meaningful and preferably intuitive way. In evolutionary biology, phylogenetic trees have long been the practice to illustrate the relationships between organisms, and when evolution is, itself, bifurcating, even very large bifurcating, phylogenetic trees are easy to interpret. In molecular biology, extremely complex interacting networks of molecules are typical, and despite the formidable complexity, a network graph can provide an intuitively accessible representation. In such a model, biological behavior is explained by information that flows through the interaction network, and part of the problem of explaining a phenotype is to identify that part of the global network that is implied by experimental data, such as differential gene expression.

Gene expression analysis and other molecular measurements performed to identify molecular correlates with changes in phenotype can identify parts of the global interaction network that control the phenotype. This is achieved by enrichment analysis against a global protein-protein and transcription factor-transcribed gene network which identifies the part of the network associated with the phenotype. In so doing, one identifies other players underlying the phenotype that may not themselves change, or may not comprise part of the data, but may nevertheless be important. For example, measurements of transcript abundance are silent with respect to possible kinase targets, but such silent nodes might be drugable.

GIRN decorates the relevant sub-network with annotation terms comprising gene ontology terms, pathway descriptors, and diseases, and does so, if one wishes, according to the same enrichment strategy used to identify the sub-network. Further, GIRN can decorate a network with drugs known to inhibit or otherwise interact with the genes in the network. This can serve as a summary skeleton for a narrative description of the phenotype suggested by the input set of genes, and also as a generator of hypotheses, by tracing network connections between drugs, genes, and pathway or ontology nodes, and trying to predict the effect of the drug on manifestation of the latter two phenotype annotations.

## Methods

### Programming

The graphs are force directed graphs implemented using the d3.js JavaScript library (www.d3js.org), jquery, html, php, and mysql. Data are stored in a mySQL database.

### Graphic conventions

An objective was to make these networks easily visually interpretable. Therefore, where possible, icons were chosen to provide a visual clue as to the class of object. For example, gene targets of a transcription factor are represented by a short double helix, drugs are represented by the standard Rx symbol, and diseases are represented by a stumbling person. Arrows reflect the order of objects, left to right, in the formatted list of relationships, and can be added or removed at will. For protein-protein interactions, this has no intrinsic meaning, but a meaning can be built in if so desired. An attempt was made to use sufficient area to display all nodes without too much overlap to read their names.

### Z-score

The sub-network of genes implied by the input list is selected by setting a minimum z-score, calculated using a binomial proportions test (24).

### Annotation term enrichment

Annotation term enrichment is taken as inversely proportional to a p-value derived from Fisher's exact test, which accounts for the number of edges connected to each gene within a GO category, pathway, or disease and the probability that the presented gene would manifest an interaction with the gene at random. If a set of genes interacts with a GO category, pathway, or disease more frequently than expected at random, this would be evidence for enrichment. Enrichment is an important concept when trying to understand biological responses to stimuli. However, differential response to stimuli does not always involve substantial differential expression of genes that influence a biological phenotype. For example, if a key regulatory gene signals via changes in the phosphorylation state of its protein, it may be important to the phenotype, but not change in expression. Therefore, p-values related to enrichment may not be relevant to certain network-interpretive goals. In particular, if the goal is to identify a drug target to disrupt a phenotype, the desired target may be statistically unremarkable.

### Technical limitations

This website has limitations to the total amount of information that can be passed from server to client and to the amount of data that can be manipulated by the server in a single request. Such strategies as attempting to visualize the entire network by listing all genes in the “Genes to Map” box are not allowed. It is not generally useful to visually peruse a network having more than a few hundred nodes.

### Global network

The global network is from CPDB (Consensus Path DataBase) (11) (version 31; downloaded 3/21/16; 200,000 interactions) and Pathway Commons (14) (http://www.pathwaycommons.org; Pathway Commons.7.All.EXTENDED_BINARY_SIF.hgnc.sif; downloaded 3/21/16; 914,165 interactions), for the Protein-Protein Interaction component, and HTRIdb (Human Transcriptional Regulation Interactions database; downloaded 3/21/16; 52,467 interactions) (5) for the Transcription Factor-Transcribed Gene component. Only “interacts-with” entries in Pathway Commons were used. All of these databases draw from smaller collections. Non-redundant union of the three lists yielded 1,145,437 interactions.

### Gene ontology

The gene ontology database used herein is from the Gene Ontology Annotation (UniProt-GOA) Database and the Gene Ontology Consortium (www.geneontology.org )(1, 2). Specifically, we used the gene_association.goa_ref_human subset (5.11.2016 release) of the UniProt-GOA database.

### Pathways

Pathway data were compiled by Kamburov et al. (11, 21-23) and include BioCarta (http://cgap.nci.nih.gov/Pathways/BioCarta_Pathways), EHMN, HumanCyc (http://humancyc.org), INOH, KEGG (http://www.genome.jp/kegg), NetPath (http://www.netpath.org), PharmGKB (http://www.pharmgkb.org), PID (http://pid.nci.nih.gov), Reactome (http://reactome.org), Signalink (http://signalink.org), SMPDB (http://www.smpdb.ca), and Wikipathways (http://www.wikipathways.org).

